# An atomistic perspective on ADCC quenching by core-fucosylation of IgG1 Fc N-glycans from enhanced sampling molecular dynamics

**DOI:** 10.1101/701896

**Authors:** Aoife Harbison, Elisa Fadda

## Abstract

The immunoglobulin type G (IgG) Fc N-glycans are known to modulate the interaction with membrane-bound Fc γ receptors (FcγRs), fine-tuning the antibody’s effector function in a sequence-dependent manner. Particularly interesting in this respect are the roles of galactosylation, which levels are linked to autoimmune conditions and aging, of core fucosylation, which is known to reduce significantly the antibody-dependent cellular cytotoxicity (ADCC), and of sialylation, which also reduces ADCC but only in the context of core-fucosylation. In this work we provide an atomistic level perspective through enhanced sampling computer simulations, based on replica exchange molecular dynamics (REMD), to understand the molecular determinants linking the Fc N-glycans sequence to the observed IgG1 function. Our results indicate that the two symmetrically opposed N-glycans interact extensively through their core trimannose residues. At room temperature the terminal galactose on the α(1-6) arm is restrained to the protein through a network of interactions that keep the arm outstretched, meanwhile the α(1-3) arm extends towards the solvent where a terminal sialic acid remains fully accessible. We also find that the presence of core fucose interferes with the extended sialylated α(1-3) arm, altering its conformational propensity and as a consequence of steric hindrance, significantly enhancing the Fc dynamics. Furthermore, structural analysis shows that the core fucose position within the Fc core obstructs the access of N162 glycosylated FcγRs very much like a “door-stop”, potentially decreasing the IgG/FcγR binding free energy. All of these factors could represent important clues to understand at the molecular level the dramatic reduction of ADCC as a result of core fucosylation and provide an atomistic level-of-detail framework for the design of high potency IgG1 Fc N-glycoforms.

## Introduction

Immunoglobulins type G (IgGs) are the most abundant antibodies in human serum(Vidarsson, G., Dekkers, G., et al. 2014). Their ability to trigger an effective immune response is dependent on their interaction with the fragment crystallizable (Fc) γ receptors (FcγRs) bound to the outer membrane of immune system effector cells(Nimmerjahn, F. and Ravetch, J.V. 2005). The interaction between IgGs and FcγRs type III (FcγRIII or CD16) specifically triggers an antibody-dependent cell-based cytotoxicity (ADCC) response(Battella, S., Cox, M.C., et al. 2016) that leads to the destruction of a targeted cell. Because ADCC is considered as the main antibody-based mechanism against tumour cells, strategies aimed at regulating ADCC are highly sought after, specifically within the framework of the antibody engineering of cancer therapeutics(Mimura, Y., Katoh, T., et al. 2018). The IgGs/FcγRs binding specificity and affinity depend not only on the specific amino acids in the IgG1 CH2 region in direct contact with the FcγR (Roberts, J.T. and Barb, A.W. 2018), but also on the type of glycosylation on both, the IgG1 Fc and the FcγR(Hayes, J.M., Cosgrave, E.F., et al. 2014, Hayes, J.M., Frostell, A., et al. 2017, Subedi, G.P. and Barb, A.W. 2018). More specifically, both CH2 domains of human IgGs are glycosylated at Asn 297 with complex biantennary N-glycans, 96% of which are core-fucosylated and 60% terminating with one or two galactose residues (Pucić, M., Knezević, A., et al. 2011). Around 20% of the Fc N-glycans are sialylated(Pucić, M., Knezević, A., et al. 2011) with one terminal sialic acid preferentially on the α(1-3) arm(Barb, A.W., Brady, E.K., et al. 2009). Decreasing IgG1 Fc galactosylation levels have been linked to autoimmune conditions, such as rheumatoid arthritis(Gudelj, I., Lauc, G., et al. 2018), and both galactosylation and especially sialylation are known to trigger an anti-inflammatory response(Pagan, J.D., Kitaoka, M., et al. 2018). The mechanism(s) linking the Fc N-glycans sequence to IgG1 function are unclear. Nevertheless, it is very likely that the molecular basis for these effects resides in how the Fc N-glycoforms modulate the IgG/FcγRs recognition and binding affinity.

The crystal structure of a complex between an IgG1 and FcγRIII gives important insight into this matter, showing that the Fc and FcγRIII N-glycans form an intricate network of interactions upon binding(Ferrara, C., Grau, S., et al. 2011). These carbohydrate-carbohydrate contacts are important in regulating the binding affinity of the complex(Ferrara, C., Grau, S., et al. 2011, Ferrara, C., Stuart, F., et al. 2006, Subedi, G.P. and Barb, A.W. 2018). Indeed, the comparison between two crystal structures of the IgG1/FcγRIII complex, one where the IgG1 N-glycan is core-fucosylated and the other where it is not(Ferrara, C., Grau, S., et al. 2011), shows that core-fucose hinders the carbohydrate-carbohydrate interaction, displacing the Fc N-glycans by of 2.6 Å in comparison to the structure with the non-fucosylated Fc-linked N-glycan(Ferrara, C., Grau, S., et al. 2011). Because of this steric hindrance, a less effective binding network can be formed between the two glycans, with a consequent reduction of the complex binding affinity. Although the effectiveness of the glycans interaction is most likely a contributing factor to the reduction of the ADCC by core-fucosylation, carbohydrate-carbohydrate binding affinities are known to be very weak(Lai, C.H., Hütter, J., et al. 2016, Spillmann, D. and Burger, M.M. 1996), thus a slight change in the enthalpic contribution due to looser N-glycans contacts is rather difficult to reconcile with the 100-fold ADCC reduction observed in the presence of a core-fucosylated Fc N-glycan (Iida, S., Misaka, H., et al. 2006, Kiyoshi, M., Caaveiro, J.M., et al. 2015, Li, T., DiLillo, D.J., et al. 2017, Shields, R.L., Lai, J., et al. 2002). Furthermore, it does not explain how sialylation can decrease ADCC but only in the context of core-fucosylation(Li, T., DiLillo, D.J., et al. 2017).

To gain further insight in the role of the Fc N-glycans sequence, structure and dynamics in the IgGs function, here we present the results of a molecular dynamics study of IgG1 Fc domains with specific N-glycans, shown in **Figure 1**, known to be significantly populated in human IgG1s(Pucić, M., Knezević, A., et al. 2011). Extensive sampling through temperature replica exchange molecular dynamics (REMD)(Sugita, Y. and Okamoto, Y. 1999) was chosen as the method to explore exhaustively the potentially rugged conformational space. The N-glycoforms were chosen specifically to address the three following points, 1) to investigate a potential link between the structure and dynamics of core-fucosylated N-glycans to the dramatic ADCC reduction, 2) to determine how the preferential conformation and dynamics of a sialylated α(1-3) arm may affect ADCC in context of core-fucosylation(Li, T., DiLillo, D.J., et al. 2017) and finally 3) to analyse the dynamics of the galactosylated α(1-6) arm in Fc-linked relative to the free N-glycans.

**Figure 1.**
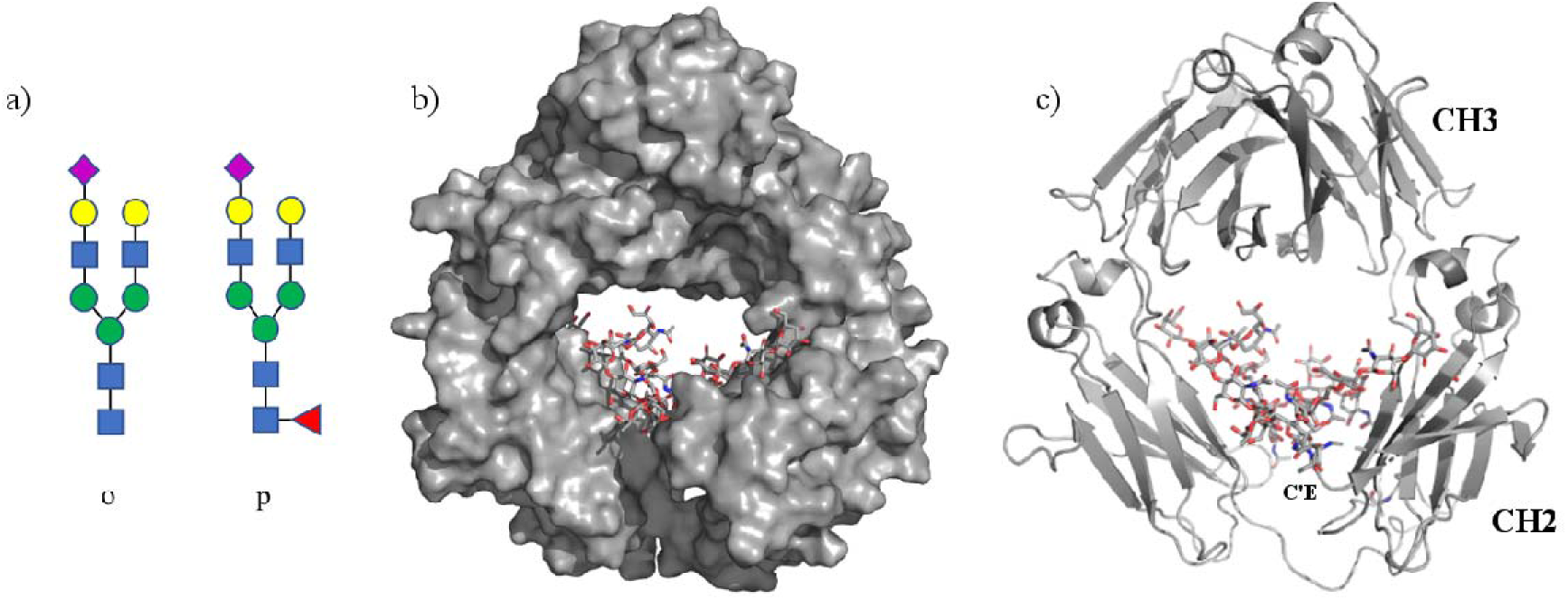
The two different types of biantennary complex N-glycans we have considered in this study are shown in panel a). For simplicity and consistently with our previous work on the unlinked form of these sugars(Harbison, A.M., Brosnan, L.P., et al. 2019), we named the core-fucosylated N-glycan sugar *p* and the non core-fucosylated N-glycan sugar *o*. A representative structure from one of the REMD simulations of the IgG1 Fc with the linked N-glycans is shown in panel b) with the protein represented as solvent accessible surface area and in panel c) with cartoon rendering, where the CH3 and CH2 domains are labelled, together with the CH2 C’E loop carrying the N297-linked N-glycan.

The simulations results show that core-fucosylation affects dramatically the dynamics of the α(1-3) arm of the opposed N-glycan and as a consequence of steric hindrance it significantly enhances the dynamics of the Fc domain. Also, structural alignment of our systems to available crystallographic data show that the core-fucose position within the Fc core may obstruct the entry of N162 N-glycosylated FcγRs, thus binding a core-fucosylated IgG1 may require a more complex conformational displacement of the CH2 domain, consistent with a higher energetic cost and a lower binding affinity. Furthermore, we find that at room temperature (300 K) the terminal galactose in the α(1-6) arm is heavily restrained to the CH2 domain of the protein, which promotes an outstretched conformation as the only significantly populated conformer. In the following sections we will present and discuss these results in detail within the framework of the available experimental data and of the known evidence of the N-glycosylation dependence of IgG1 effector functions.

## Computational Method

Protein and carbohydrate atoms were represented with the ff14SB(Maier, J.A., Martinez, C., et al. 2015) and GLYCAM06j-1(Kirschner, K.N., Yongye, A.B., et al. 2008) forcefields, respectively, while the TIP3P model (JORGENSEN, W., CHANDRASEKHAR, J., et al. 1983) was used to represent the solvent. The total electrostatic charge of the system was neutralized by adding Cl^−^ ions. All simulations were carried out using NAMD version 2.31b1(Phillips, J.C., Braun, R., et al. 2005). Each system was prepared through an initial 500k cycles of conjugate gradient minimisation with a restraint of 5 kcal mol^−1^Å^−2^ on all heavy atoms. The cutoff for van der Waals interactions was set to 12 Å and smoothing functions were applied between 11 and 13.5 Å. Particle Mesh Ewald (PME) was used to treat electrostatic interactions with a charge grid of 1 Å and a sixth order spline function for mesh interpolation. All non-bonded interactions not directly connected were excluded, namely 1-3 pair interaction, or scaled by 0.8333, namely 1-4 pair interaction. Following energy minimization, the systems were heated from 0 to 300 K over 600 cycles with restraints on all heavy atoms in place. For the temperature Replica Exchange Molecular Dynamics (REMD)(Sugita, Y. and Okamoto, Y. 1999) 90 replicas were generated in the temperature range between 300 and 500 K. The systems were equilibrated with restraints on the protein backbone atoms and on the glycans heavy atoms for 500 ps, followed by 500 ps of unrestrained equilibrations for each replica. The production steps ranged between 13 ns and 11 ns for each replica, with an integrated time step of 2 fs. The SHAKE algorithm was used to restrain bonds to hydrogen atoms. The simulations were carried out on six IgG1 Fc models in total, one with two core-fucosylated N-glycans (*pp*), namely sugar *p* shown in **Figure 1**, one with two non-fucosylated N-glycans (*op*) and another with one core-fucosylated and one non-fucosylated N-glycan (*op*). Because the molecular crowding within the Fc core could limit the conformational space even within an enhanced sampling scheme, in order to explore the possibility of the folding (closing) of the galactosylated α(1-6) arm we observed for the corresponding unlinked glycans, we built all starting structures with α(1-6) arms both in the open and closed conformations, see **Figure S.1**. Data analysis and structural alignments were done with VMD v.1.9.3 beta 1(Humphrey, W., Dalke, A., et al. 1996) and *seaborn* (https://seaborn.pydata.org) was used for all statistical data visualization. Further information on the system set-up is included in the Supplementary Material.

## Results

The results below are presented in separate sections for clarity. Unless stated otherwise, all results refer to simulations started with the N-glycans α(1-6) arm in the open conformation.

### Protein and Fc-linked N-Glycans Dynamics

The IgG1 Fc domains flexibility and its overall dynamics has been evaluated in terms of backbone RMSD values, calculated from the simulations relative to the 1.9 Å resolution crystal structure (PDBid 4DZ8) of a N-glycosylated and core-fucosylated IgG1 Fc(Strop, P., Ho, W.H., et al. 2012). The structural alignment was done on the two symmetric CH3 domains alone for all simulations as it is the most stable part of the IgG1 Fc. Results are shown in **Figure 2** in terms of kernel density estimation (KDE) plots of the protein backbone RMDS values obtained from simulations of an IgG1 Fc linked to two core-fucosylated sugar *p* on both sides (*pp*), see **Figure 2 panel a**, an IgG1 Fc linked to two non-fucosylated sugar *o* on both sides (*oo*), see **Figure 2 panel b**, and to an IgG1 Fc with one sugar *o* and one sugar *p* on each side (*op*), see **Figure 2 panels c** and **d**. The average backbone RMSD values for all the systems studied are shown in **Table S.1**. The results indicate that core-fucosylation significantly enhances the protein dynamics, with a maximum effect when both Fc N-glycans are fucosylated, see **Figure 2 panel a**. As it will be discussed in detail further below, this enhanced dynamic is a consequence of the steric interaction between the core-fucose of one N-glycan and the α(1-3) arm of the symmetrically opposed N-glycan. The KDE distributions also indicate that the protein dynamics is primarily determined by the CH2 domain, which is linked to the more rigid CH3 domain by an unstructured loop, see **Figure 1 panel c**. that allows for a flexible architecture. The stability of the CH3 domain is not affected by the presence of core-fucose, nor by the conformational dynamics of the arms. The results obtained for the *op* IgG1 Fc show that the presence of even one core-fucose increases the dynamics of the CH2 domain, even if to a lesser degree than two core-fucosylated N-glycans. The CH2 linked to the non-fucosylated sugar *o*, see **Figure 2 panel c**, appears slightly more dynamic, as a consequence of the steric hindrance between sugar *o*’s α(1-3) arm and the core-fucose on the facing sugar *p*.

**Figure 2.**
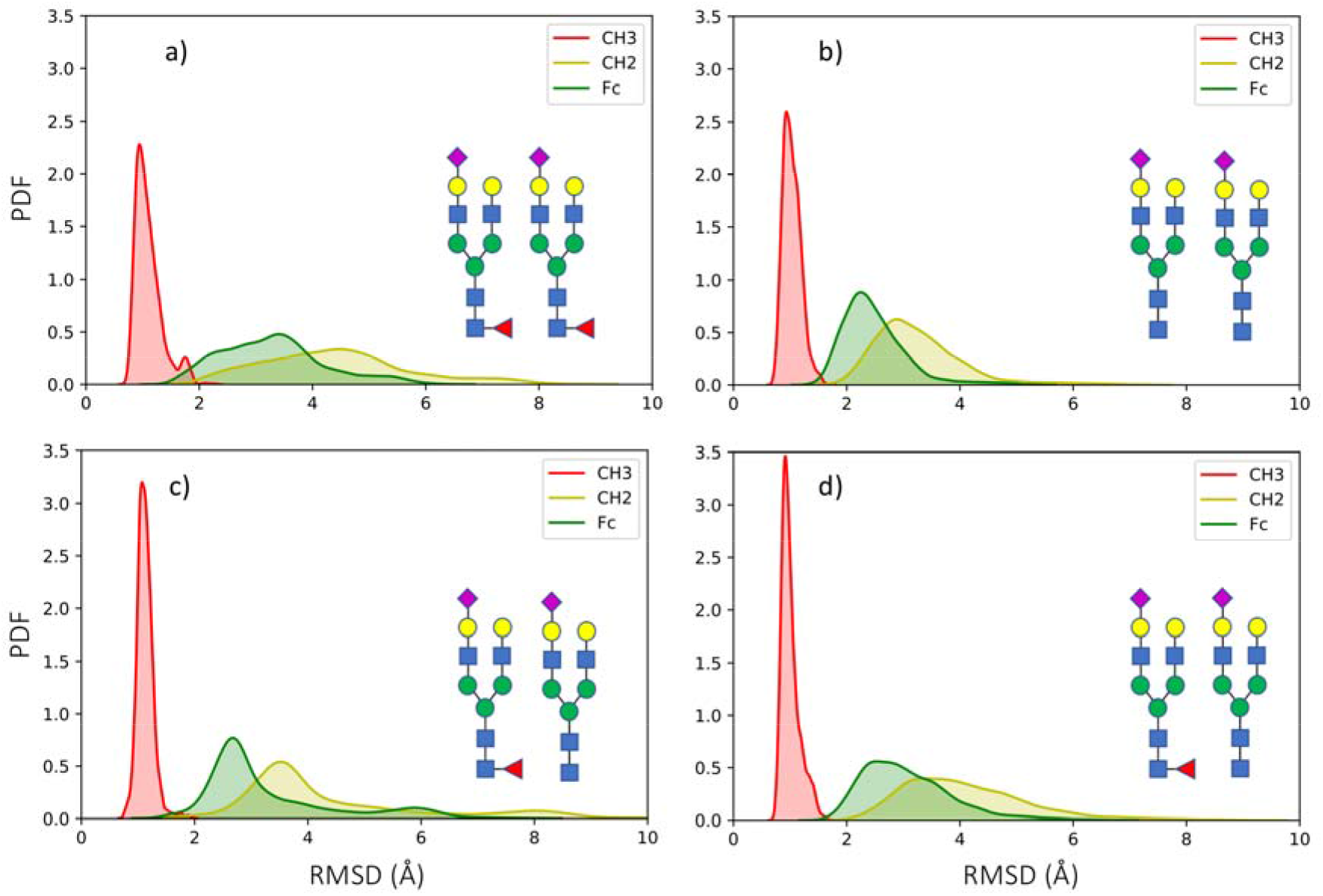
KDE plots of protein backbone RMSD values calculated throughout the REMD simulation started from open conformations of the α(1-6) arms on an IgG1 Fc with two core-fucosylated sugars *p* in panel a) with two non core-fucosylated sugar *o* in panel b) and with one core-fucosylated sugar *p* and one non core-fucosylated sugar *o* in panels c) and d), where the two CH3-CH2 domains are represented separately for the non core-fucosylated and for the core fucosylated Fc-linked CH2 in panels c) and d), respectively. The probability density function (PDF) is on the y axis and the root mean square deviation (RMSD) values on the x axis. The plots are. Based on 6500 points for panel a), and 5500 points for panels b), c) and d).

In regards to the dynamics α(1-6) and α(1-3) arms, the RMSD values distributions are shown in **Figure 3** for the *pp, oo* and *op* IgG1 Fcs. The distributions show that the presence of core-fucosylation enhances the dynamics of the whole N-glycans, but especially of the α(1-3) arms. This enhanced flexibility is due to the steric hindrance between the core-fucose of one N-glycan and the extended sialylated α(1-3) arm of the other. This clash pushes the α(1-3) arm to interconvert between an extended conformation, which requires the opening of the CH2 domains in order to fit, and a bent conformation, where the sialic acid points towards the core of the Fc instead of towards the bulk water, see **Figure 4**. The extended α(1-3) arm is the preferential conformation seen during the simulation of the *oo* system and corresponds to the maxima in **Figure 3 panel d**, where none of the glycans are core-fucosylated. As a note, the shift in the RMSD values observed for glycan 1 (g1) in **Figure 3 panel d** (red line) depends on the fact that the RMSD values are calculated relative to the starting conformation, which in this specific case was a low populated one where the α(1-3) arm is slightly bent interacting with the opposite N-glycan chitobiose core. The results shown in **Figure 3 panels e** and **f**, obtained for the mixed *op* IgG1 Fc system are quite interesting in that they show that the α(1-3) arm of the non-fucosylated glycan g2 is more dynamic than the α(1-3) arm of the fucosylated g1, as it does interact with the g1 core-fucose. Also, the dynamics of α(1-6) arm of the core-fucosylated glycan g1 is directly affected by this interaction. The corresponding average RMSD values calculated for the N-glycans heavy atoms are shown in **Table S.2**.

**Figure 3.**
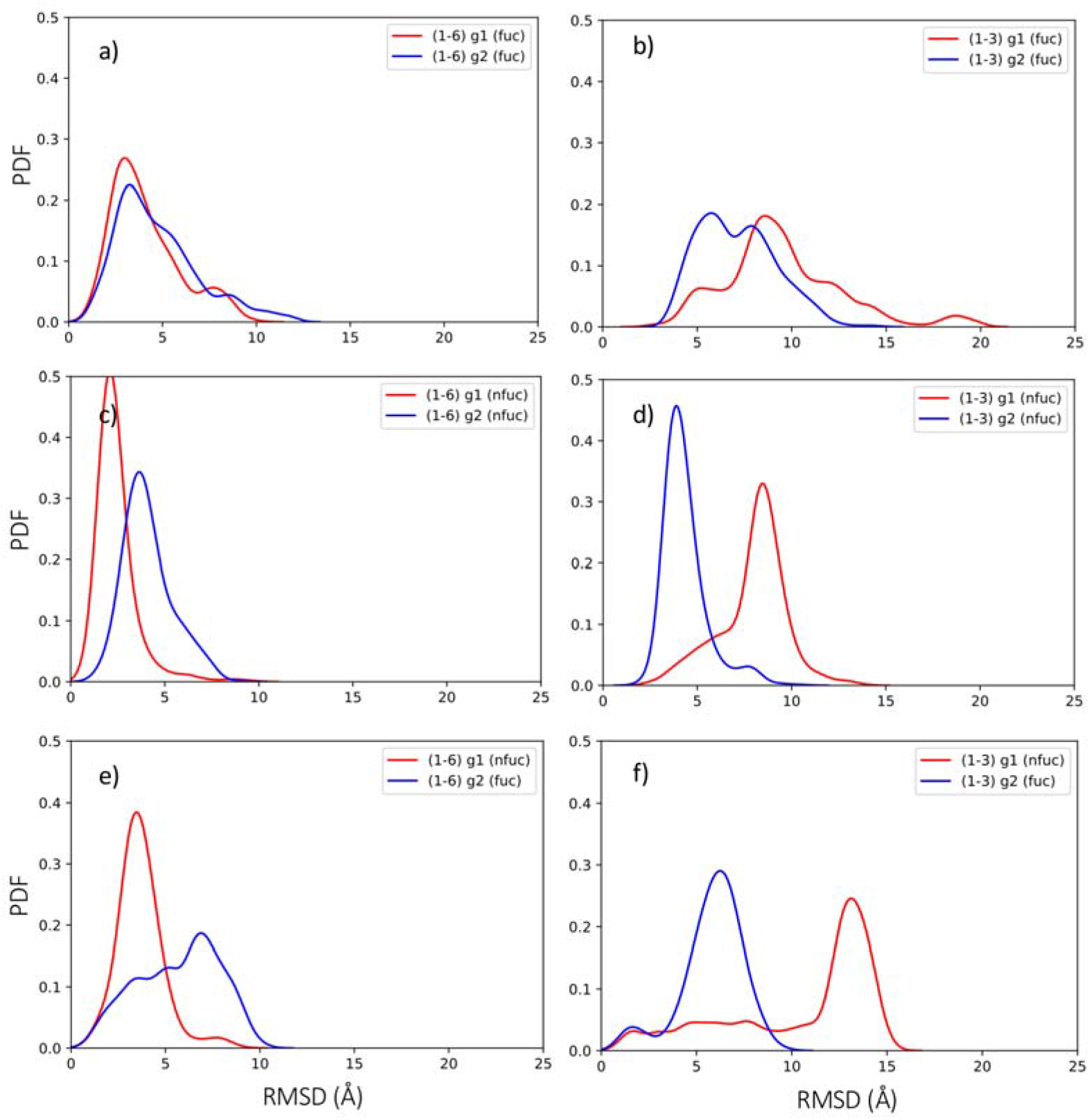
KDE plots of the RMSD values calculated for the a(1-6) and a(1-3) arms (heavy atoms) during the simulations of the IgG1 Fc with two fucosylated N-glycans (*pp*) in panels a) and b), and with two non-fucosylated N-glycans (*oo*) in panels c) and d) and with one fucosylated and one non-fucosylated N-glycans (*op*) in panels e) and f). The red and blue lines refer to the two different glycans g1 and g2, respectively.

**Figure 4.**
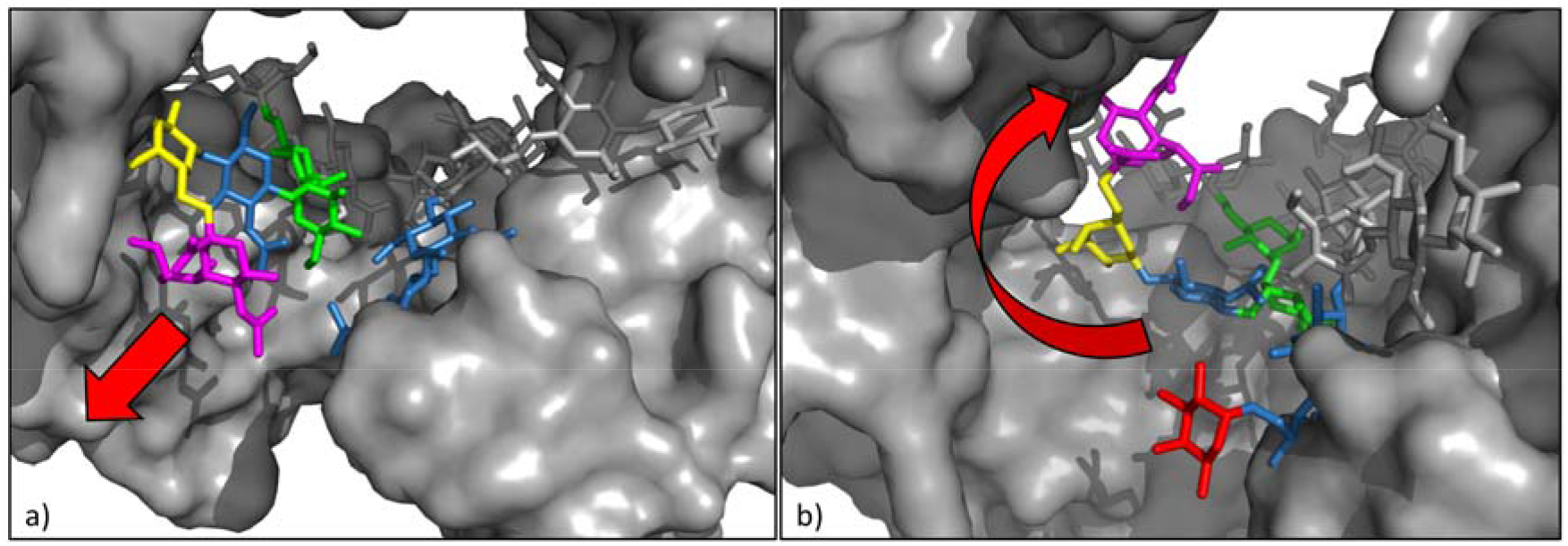
Different orientations of the sialylated α(1-3) arm in the absence and presence of core-fucose on the opposite N-glycan. In panel a) the snapshot from the *oo* IgG1 Fc simulation shows the α(1-3) arm outstretched towards the solvent, meanwhile in panel b) the snapshot from the *pp* IgG1 Fc simulation shows the α(1-3) arm folded over the Fc core obstructed by the core-fucose of the opposite N-glycan. The important residues of the N-glycans are highlighted according to the SNFG nomenclature, the protein is represented through a solvent accessible surface in grey. Rendering was done with *pyMol*.

The folding backwards of the α(1-3) arm towards the inside of the Fc core promoted by the core-fucose of the opposite glycan does not involve changes from the equilibrium torsion angle values we have determined for the unlinked (free) N-glycans in earlier work(Harbison, A.M., Brosnan, L.P., et al. 2019). As shown in **Table S.3** and **Figure S.2** the relative populations and values of the torsion angles for the α(1-3) arm are quite similar to the ones we determined in solution(Harbison, A.M., Brosnan, L.P., et al. 2019). The only significant difference is the increase of the population of the α(1-3) ψ = 96° (15) torsion to 55%, corresponding to the value in solution ψ = 102° (14) at 39%(Harbison, A.M., Brosnan, L.P., et al. 2019), which determines the bent conformation, see **Table S.3**.

### N-Glycans interactions within the Fc core

In all simulation started from an open conformation, the α(1-6) arm remains outstretched over the CH2 β sheet and restrained to it through an extensive and organized architecture of interactions with both hydrophobic and hydrophilic residues. We did not observe any unbinding events at room temperature (300 K). As shown in **Figure 5**, residues Lys 240 can interact with the carbonyl oxygen of the α(1-6) GlcNAc, meanwhile Glu 252, Asp 243 and Thr 254 are all engaging with the α(1-6) terminal Gal. Residues Phe 237, Val 256, Val 297 and Val 299 line-up to form and hydrophobic patch that supports by stacking the α(1-6) arm movement across the CH2 domain. This outstretched conformation is the only one populated in all simulations started from an open α(1-6) arm conformation, as shown by the heat map in **Figure 5**. As explained in the Method section, all simulations were also started from a closed α(1-6) arm conformation, see **Figure S.1**, where the α(1-6) arm is folded over the chitobiose core as seen for the corresponding N-glycans in solution(Harbison, A.M., Brosnan, L.P., et al. 2019). Because of the molecular crowding and the limited space available due to the partial collapse of the Fc core, in none of these simulations the N-glycan can open.

**Figure 5.**
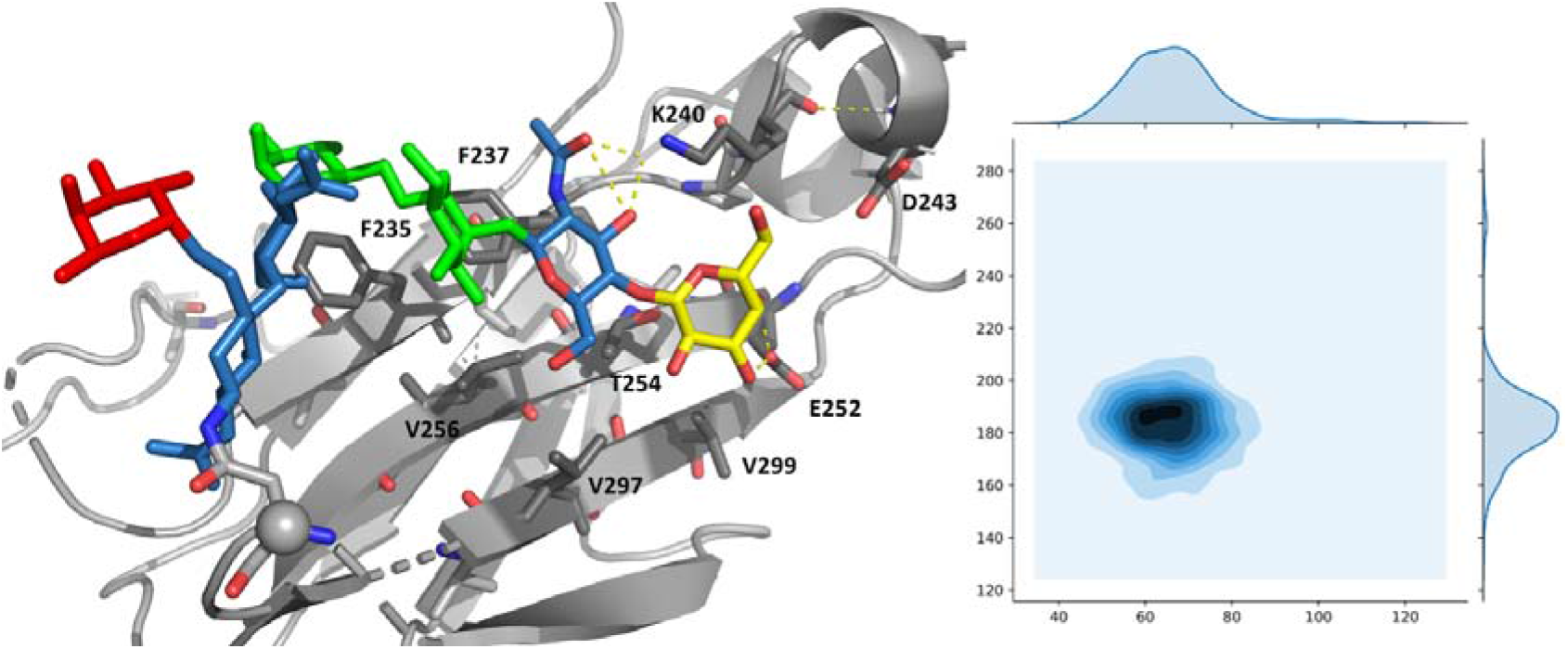
On the left-hand side, a representative image of the highest populated conformer observed for the α(1-6) arm in the simulations of both core-fucosylated (sugar *p*, represented here) and also of non core-fucosylated sugar *o*. The α(1-3) arm and the rest of the protein are not represented for clarity and the residues are coloured according to the SNFG nomenclature. On the right-hand side, the corresponding phi/psi Ramachandran plot of the α(1-6) torsion calculated from the REMD simulation of the *pp* IgG1 Fc system, showing that only the extended conformer is populated at 300 K.

The two N-glycans interact quite extensively with each other through transient and interchanging hydrogen bonds. This interaction network involves primarily the trimannose residues, while the arms are not primarily involved. A representative structure from the simulation of the *oo* IgG1 Fc is shown in **Figure S.3**, where is also evident the narrow space between the α(1-3) arm of one N-glycan and the CH2-linked GlcNAc of the other, which leaves very little room for the α(1-6) fucose.

### Core-fucosylation hinders FcγR access to the binding site

To evaluate a potential direct effect of the core-fucose in the binding of the FcγRs we performed different structural alignments of representative conformations obtained throughout our simulations to crystal structures of IgGs Fcs in complex with FcγRs, one with N-glycosylated FcγRIII at Asn 162 with PDBids 3SGJ and 3SGK(Ferrara, C., Grau, S., et al. 2011) and one where the FcγRIII is non-glycosylated with PDBid 1E4K (Sondermann, P., Huber, R., et al. 2000). As shown in **Figure 6 panel a**, the alignment of our structure to the 3GSJ based exclusively on the protein residues of the CH2 and CH3 domains (backbone RMSD of 1.9 Å based on 196 atoms) on the left-hand side of the Figure, shows that the accommodation of the FcγRIII requires a significant displacement of the opposite CH2 domain. This is regardless of the core-fucosylation state of the Fc N-glycan linked to that specific CH2 domain. The obstructing effect of the core-fucose is quite evident when we align the C’E loops, i.e. from residue 293 to 300, to identify the relative position of the Fc N-glycan relative to the FcγRIII residues and to the N-glycan at Asn 162 on the FcγRIII when sufficient space has been created to accommodate the FcγRIII due to the CH2 displacement. As shown in **Figure 6 panel b**, the position of the core-fucose does indeed obstruct the access of the FcγRIII due to a steric clash with the N-glycan at Asn 162. The higher energetic cost of moving the fucosylated N-glycan in addition to the CH2 domain displacement is consistent with a lower binding affinity of the complex. The alignment of our core-fucosylated *pp* IgG Fc to the structure of a complex with FcγRIII (PDBid 1E4K)(Sondermann, P., Huber, R., et al. 2000) shows that in the absence of the N-glycan at Asn 162 the fucose is not hindering the binding, see **Figure S.4**.

Nevertheless, a lower binding affinity may be expected because of the missing interactions of the IgG Fc with the FcγRIII Asn 162 N-glycan(Subedi, G.P. and Barb, A.W. 2018).

**Figure 6.**
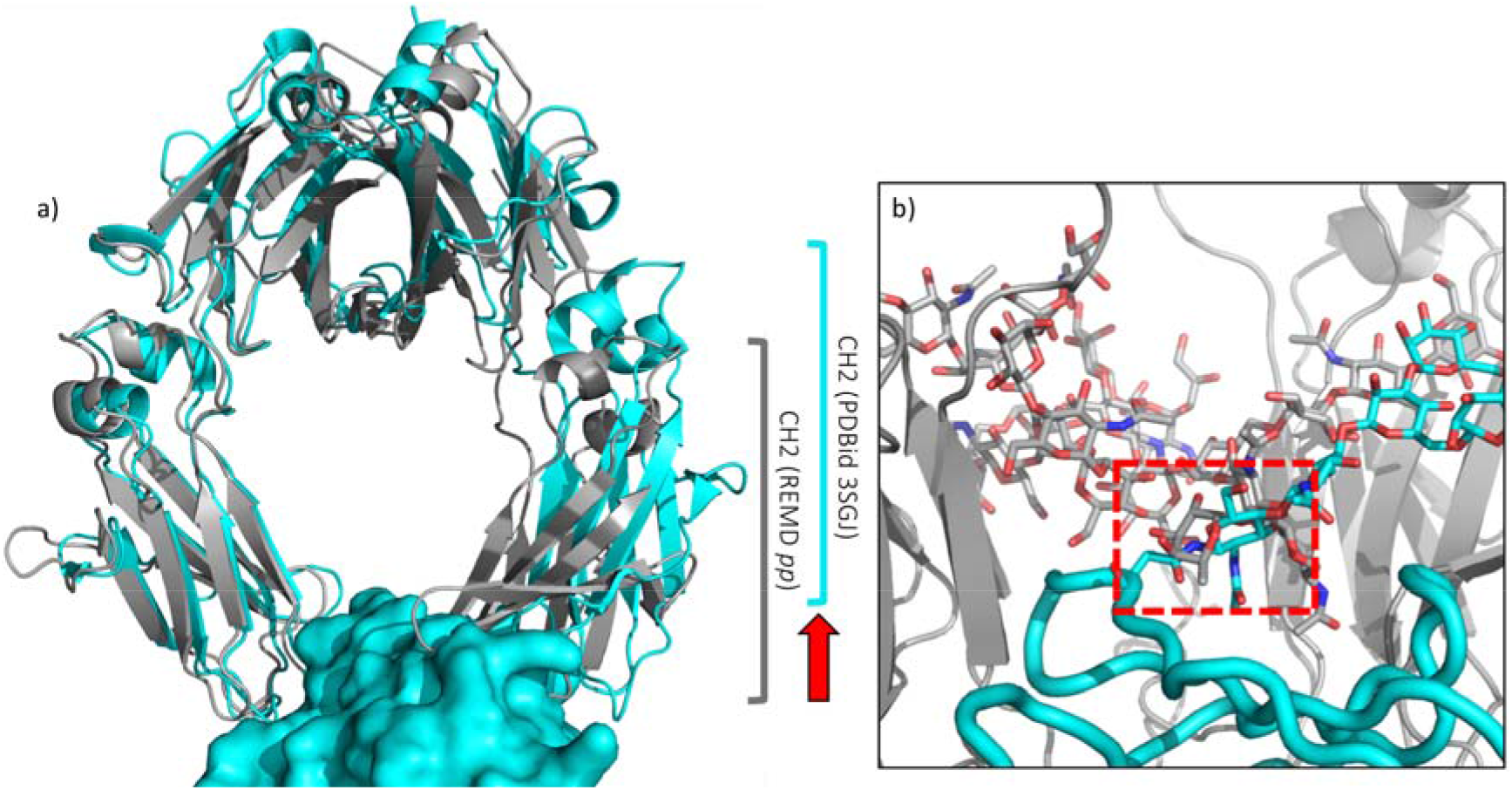
Structural alignments of a representative structure from the REMD simulation of the *pp* IgG1 Fc (grey) with the IgG Fc in complex with the FcγRIII PDBid 3SGJ (cyan). In panel a) the FcγRIII is shown as a solvent accessible surface while the IgG Fc are represented with cartoon rendering. The N-glycans are omitted for clarity purposes. The CH2 domain shift is represented through a red arrow, while the brackets span the length of the CH2 domain. In panel b) the FcγRIII backbone is represented in cyan as tube while the glycan as sticks. The structure from *pp* IgG1 Fc MD aligned through the C’E loop to the PDBid 3SGJ is represented in grey, with the Fc N-glycan also in sticks. The clash between the core fucose and the N162 N-glycan on the FcγRIII is highlighted within a red-dashed square.

## Discussion

In this work we have determined the conformational propensities of two specific N-glycans, shown in **Figure 1**, linked to the IgG1 Fc to identify how their sequence and structure modulates the dynamics of the system and in turn how can it affect the FcγRs molecular recognition and binding affinity. We designed the study specifically to address the still open question on how core-fucosylation of Fc-linked N-glycans reduces drastically the IgGs ADCC especially in the context of sialylation(Li, T., DiLillo, D.J., et al. 2017). The comparison between core-fucosylated and non-fucosylated systems shows that core-fucose affects the position of the sialylated α(1-3) arm of the opposite N-glycan because of steric hindrance. As a consequence, the α(1-3) arm becomes increasingly dynamic, switching between an outstretched and a bent conformation where the sialic acid is directed towards the Fc core. The α(1-3) arm outstretched conformation is found to be the highest populated in non-fucosylated systems. Accommodating an outstretched α(1-3) arm beside the core fucose translates into a widening of the Fc core, thus it has a significant effect in enhancing the dynamics of the Fc, in particular at the level of the CH2 domains, which are quite flexible due to the Fc architecture. The higher level of conformational disorder of the protein and the seclusion of the sialic acid from the interaction with the incoming FcγRs could both be determinants in decreasing the levels of molecular recognition and ultimately in weakening the binding affinity. Furthermore, structural alignments of representative conformers from our simulations with different crystal structures of IgG1 Fcs in complex with FcgRIIIs(Ferrara, C., Grau, S., et al. 2011, Sondermann, P., Huber, R., et al. 2000) show that core-fucose obstructs the entry of a Asn 162 N-glycosylated FcgRIII, posing a further burden in terms of the energy required to displace it in addition to the displacement of the CH2 domain. This information derived from the structural alignments of N-glycosylated IgG1 Fcs is quite helpful in that it provides for the first time a view of how the correct structure and dynamics of the Fc-linked N-glycans work within the framework of the IgG1 Fc/FcgRIIIs complex. Indeed, because of their intrinsic dynamics, in most crystal structures the N-glycans are either invisible or fitted to very unusual (and debatable) conformations.

In previous work(Harbison, A.M., Brosnan, L.P., et al. 2019) we have determined the intrinsic dynamics and conformational propensities of all N-glycans significantly populated in IgG1s when unlinked from the IgG, i.e. free in solution. We found that the galactosylation of the α(1-6) arm results in a dramatic change in the N-glycan preferential conformation, where the α(1-6) arm is folded over the chitobiose core, instead of being outstretched(Harbison, A.M., Brosnan, L.P., et al. 2019). Such compact structure is consistent with a more difficult recognition of the α(1-6) in unbound N-glycans from lectins(Echeverria, B., Serna, S., et al. 2018) and from sialyltransferases(Barb, A.W., Brady, E.K., et al. 2009), and it can also possibly explain the interdependence of the N-glycans arms functionalization process(Moginger, U., Grunewald, S., et al. 2018). Notably, the α(1-6) arm folding is independent of the presence of a terminal α(2-6) sialic acid, of the sequence on the α(1-3) arm and of core-fucosylation(Harbison, A.M., Brosnan, L.P., et al. 2019). In an Fc-linked N-glycan the terminal galactose on the α(1-6) arm is firmly restrained to the CH2 domain through a network of hydrogen bonds and hydrophobic interaction and we observed no α(1-6) unbinding nor folding events at 300 K. This is somewhat is disagreement with NMR data that suggest that the Fc-linked α(1-6) arm has a structural behaviour in between a restrained and a free (unlinked) N-glycan at 15 °C and at room temperatures(Barb, A.W. and Prestegard, J.H. 2011). Nevertheless, in agreement with the NMR study above we observe that the free N-glycan behaviour, which we assume to be the form unrestrained from the CH2 domain, increases with temperature, see **Figure S.5**. Indeed, as the temperature raises the α(1-6) arm becomes progressively more dynamic and at the extreme of 500 K is completely unrestrained. Therefore, it is possible that the strength of the hydrogen bonding interactions provided by the force field representation is too high at 300 K relative to the experimental conditions, but they get increasingly more balanced as the temperature is increased, providing the correct trend. Additionally, based on these data and analysis the reasons why the galactosylated α(1-6) arm is more difficult to functionalize both in free and in Fc-linked N-glycans are actually different. In the former case a compact folded structure preclude access relative to the more accessible and outstretched α(1-3) arm, while in the case of an Fc-linked N-glycan the strong interaction of the terminal galactose with the CH2 domain segregates the α(1-6) arm from the solvent, thus precluding accessibility.

## Conclusions

In this work we used temperature REMD to assess the role of core-fucosylation of the Fc-linked N-glycans in the IgG1 structure and interaction with FcγRs. The results indicate a significant enhancement of the dynamics of the protein and of the N-glycans intrinsic structural disorder. Additionally, we suggest a mechanistic pathway for the binding of FcγRs where core-fucose functions as a “door-stop” to the access of Asn 162 N-glycosylated FcγRs. These atomistic level, dynamic information provides for the first time to our knowledge a working framework for the rational design of IgG1 N-glycoforms that can enhance ADCC.

## Acknowledgements

EF would like to thank Dr Nina Weisser (Zymeworks, Vancouver, BC) for insightful discussions on the project. The authors gratefully acknowledge the Irish Centre for High-End Computing (ICHEC) for the generous allocation of computational resources. The John and Pat Hume Doctoral Awards programme at Maynooth University is also gratefully acknowledged for funding.

