## Supplementary Material for "An atomistic perspective on ADCC quenching by core-fucosylation of IgG1 Fc N-glycans from enhanced sampling molecular dynamics"

**Computational Method**

The crystal structure of the human Fc fragment with PDBid 1FC1 was used as a starting point for the preparation of all our systems. The N-glycans in this structure were functionalized to match our chosen glycoforms, therefore a β(1-4) galactose and α(2-6) sialic acid were added to the α(1-3) arm using the CHARMM GUI PDB reader tool(Park, S.J., Lee, J., et al. 2019). This structure was edited using the academic version of Schrödinger’s Maestro v.10.7.015 to rotate the torsion angles of the glycosidic linkages to release any steric clash that may have occurred during the build. The values of the glyosidic linkage torsion angles were taken from our earlier work on the unlinked glycans(Harbison, A.M., Brosnan, L.P., et al. 2019). Protein residues Cys 239 to Gly 250 from the human IgG B12 with PDBid 1HZH were linked to the structurally aligned 1FC1 using Maestro to build the disulfide bonds that keep the Fc region stable during the simulation. To ensure by construction that a potential folded conformation of the of the α(1-6) arm was sampled, all simulations were started with both outstretched (open) and folded (closed) conformations of the arm. Maestro(Schrodinger, Maestro, 2012) was also used to modify the α(1-6) glycosidic linkage to get the open and closed conformations for a total of six starting structures.

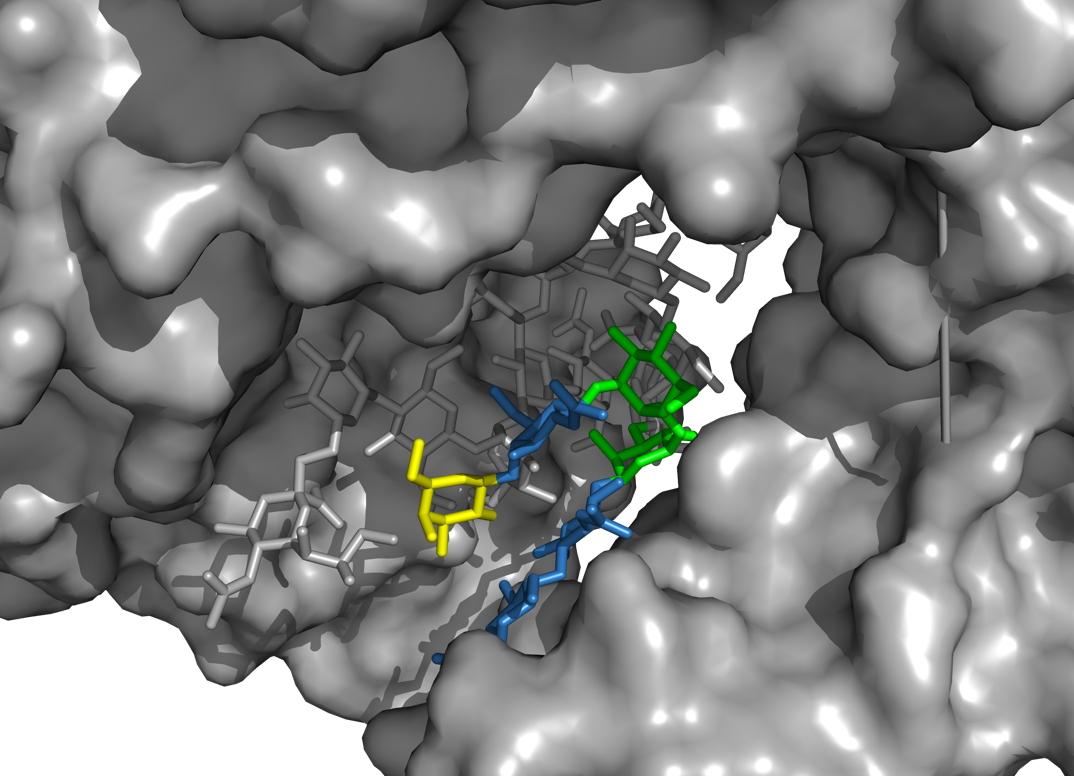

**Figure S. 1** Structure of the α(1-6) arm in a snapshot from the end of the oo IgG1 Fc closed simulation (11 ns x 90 replicas) shows an extremely compact Fc core where the N-glycans are compressed together. The chitobiose and the α(1-6) arm are highlighted with SNFG colouring, while the protein is showed through a solvent accessible area in gray.

**Table S. 1** Average protein backbone RMSD values (Å) calculated through the REMD simulations of the IgG Fc linked to different sets of N-glycans. Sugars o and p are shown in **Figure 1**. Standard deviation values are shown in parenthesis. The “open” and “closed” labels correspond to simulations started from an outstretched (open) or from a folded-over (closed) conformation of the α(1-6) arm.

|  | **Fc** | | **CH2** | | **CH3** | |
| --- | --- | --- | --- | --- | --- | --- |
| **N297-glycans** | **open** | **closed** | **open** | **closed** | **open** | **closed** |
| ***pp*** | 3.3 (0.9) | 3.3 (0.9) | 4.4 (1.3) | 4.4 (1.2) | 1.1 (0.2) | 1.0 (0.1) |
| ***oo*** | 2.5 (0.6) | 3.9 (1.3) | 3.3 (0.8) | 5.2 (1.8) | 1.0 (0.2) | 1.1 (0.2) |
| ***op (nFuc side)*** | 3.1 (0.8) | 3.4 (0.8) | 4.1 (1.1) | 4.6 (1.0) | 1.0 (0.2) | 1.1 (0.1) |
| ***op (Fuc side)*** | 3.3 (1.2) | 4.1 (1.3) | 4.4 (1.7) | 5.6 (1.8) | 1.1 (0.1) | 1.1 (0.2) |

**Table S. 2** Average RMSD values (Å) calculated over all N-glycans heavy atoms. For each N-glycan the alignment was dove over all the heavy atoms of the core chitobiose and the arms were considered from the central Man. Standard deviation values are shown in parenthesis.

|  | **α(1-6)** | | **α(1-3)** | |
| --- | --- | --- | --- | --- |
| **N297-glycans** | ***g1*** | ***g2*** | ***g1*** | ***g2*** |
| ***pp*** | 4.1 (1.8) | 4.7 (2.2) | 9.5 (3.2) | 7.1 (2.0) |
| ***oo*** | 2.4 (1.1) | 4.1 (1.3) | 8.0 (1.8) | 4.4 (1.1) |
| ***op*** | 3.7 (1.9) -*nf-* | 5.7 (2.1) -*f-* | 10.4 (3.9) -*nf-* | 5.8 (1.6) -*f-* |

**Table S. 3** Averaged torsion angle values calculated for one of the two α(1-3) arms (g1) during the simulation of the pp IgG Fc. Standard deviations are indicated in parenthesis and populations (%) are highlighted in red.

| ***pp* α(1-3)** | **phi** | | **psi** | | **omega** | |
| --- | --- | --- | --- | --- | --- | --- |
| Man-α(1-3)-Man | 68 (9) 100 | - | 96 (15) 55 | 138 (11) 45 | - | - |
| GlcNAc-β(1-2)-Man | -77 (12) 82 | -116 (14) 18 | 178 (44) 79 | 98 (13) 21 | - | - |
| Gal-β(1-4)-GlcNAc | -72 (13) 95 | 61 (11) 5 | -118 (15) 100 | - | - | - |
| Sia-α(2-6)-Gal | 66 (11) 93 | -50 (16) 7 | -178 (27) 87 | 97 (17) 11 | -177 (15) 44 | -62 (19) 38 |

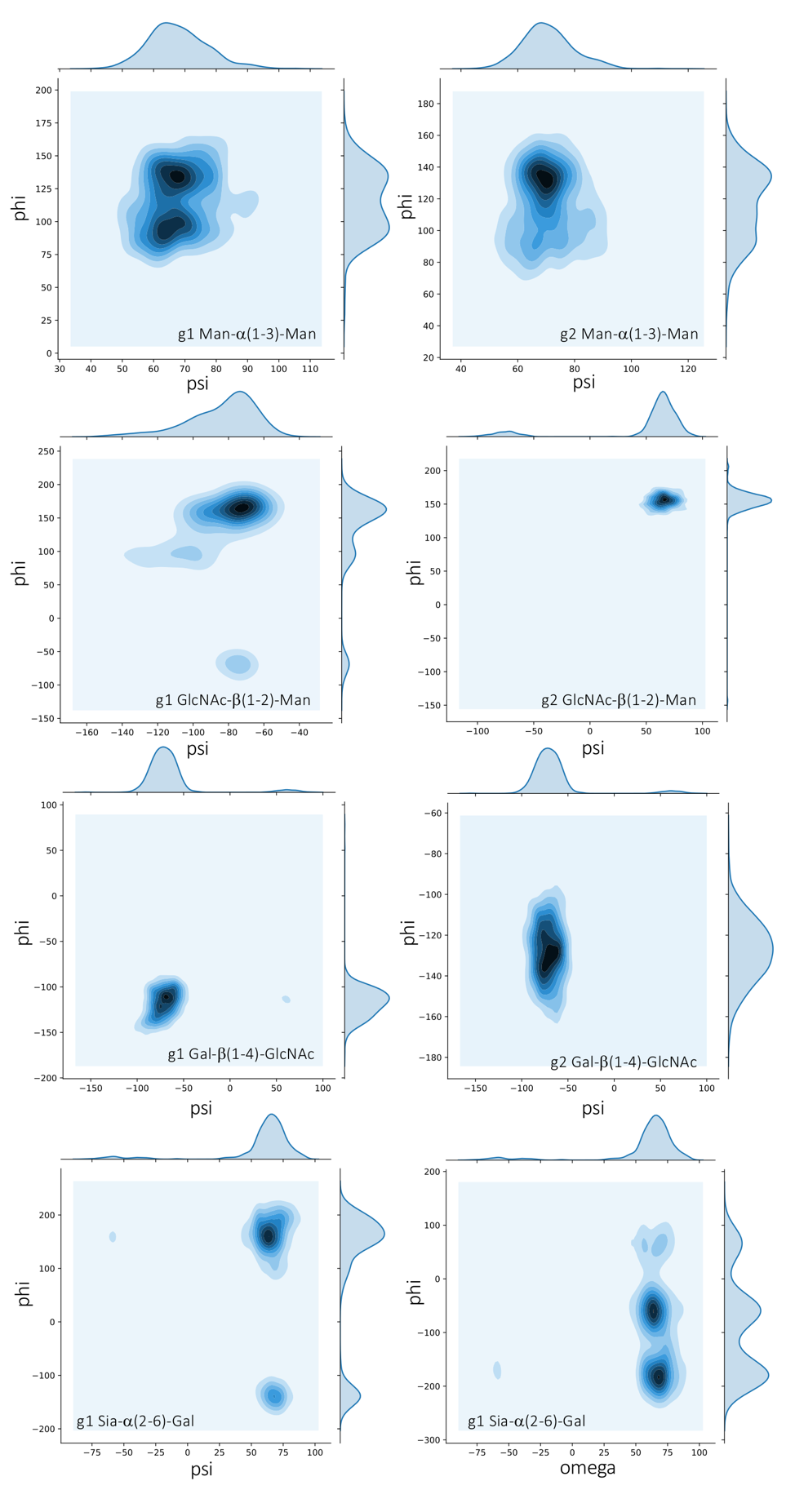

**Figure S. 2** Heat maps (Ramachandran plots) representing the conformational propensities in terms of torsion angles of the α(1-3) arms throughout the simulation of the pp IgG Fc. Only the data on g1 are shown for the Sia-α(2-6)-Gal linkage for simplicity. The maps are built on 6500 data points per axis and were made with seaborn.

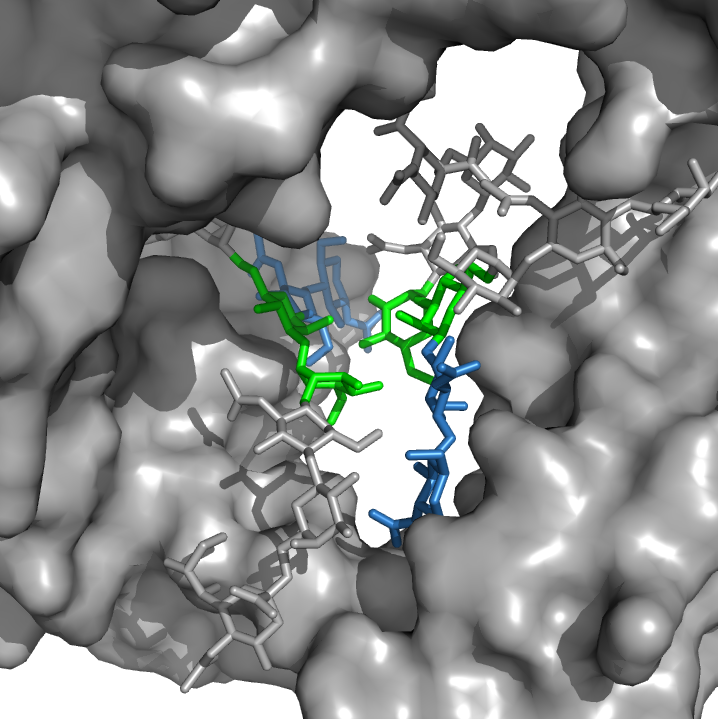

**Figure S. 3** The two trimannose cores (highlighted by SNFG colouring) in the symmetrically opposed N-glycans shown in a snapshot from the simulation of the oo IgG1 Fc. The interactions between the residues are primarily hydrogen bonds and are interchanging continuously throughout the simulation.

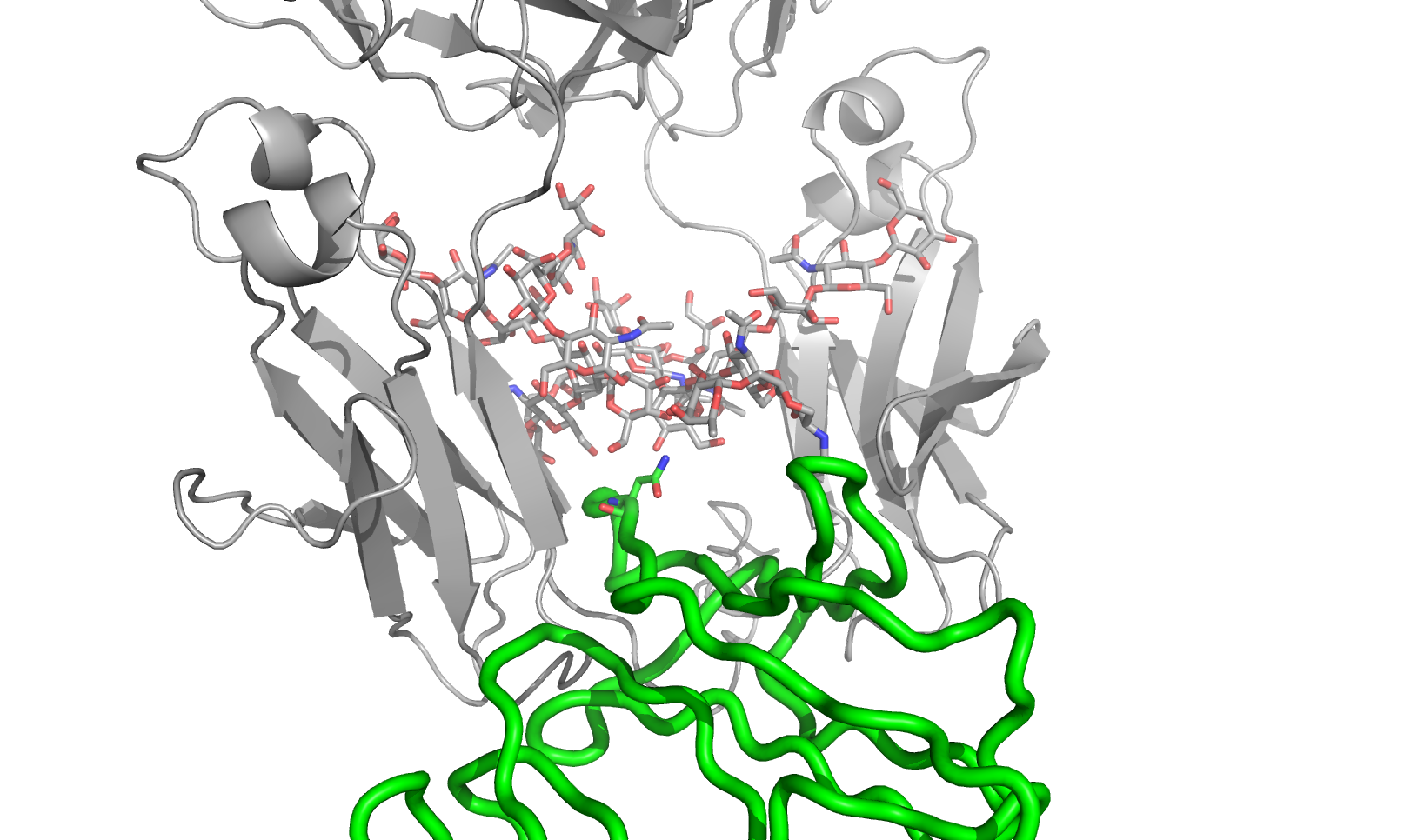

**Figure S. 4** Alignment of a representative structure from the pp IgG1 Fc REMD simulation represented in grey and the structure of the complex between the IgG1 Fc and the FcγRIII, shown in green tubes, with PDBid 1E4K(Sondermann, P., Huber, R., et al. 2000). The IgG1 Fc from PDBid 1E4K is omitted for clarity. The position of the core fucose is highlighted in a red-dashed frame, the Asn 162 of the FcγRIII is represented with sticks for clarity.

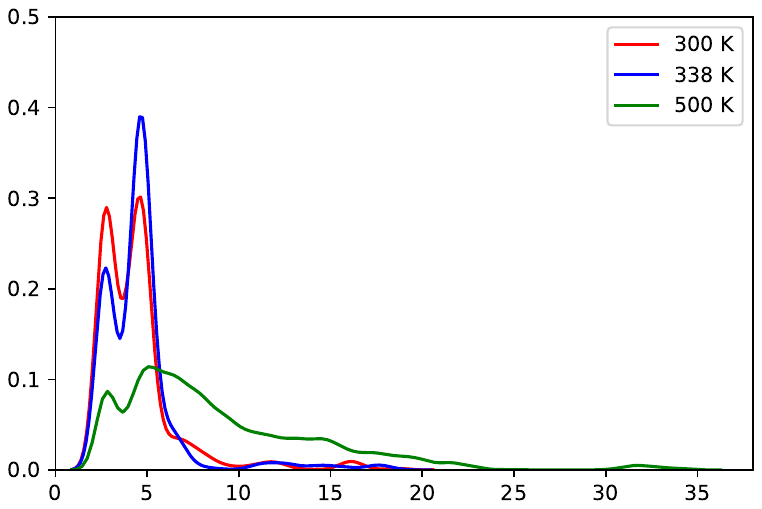

RMSD (Å)

PDF

**Figure S. 5** KDE distributions of the distance (Å) between one of the CH2 Glu 252 carboxylic oxygen and the O4 of the terminal Gal in the N-glycan α(1-6) arm in function of temperature. This distance was used as a parameter to gauge the position of the α(1-6) arm relative to CH2 and to identify the bound (values within hydrogen bond distance) from the unbound (values above hydrogen bond distance).

1.
